# Structural characterization of cocktail-like targeting polysaccharides from *Ecklonia kurome* Okam and their anti-SARS-CoV-2 activities *invitro*

**DOI:** 10.1101/2021.01.14.426521

**Authors:** Shihai Zhang, Rongjuan Pei, Meixia Li, Hao Sun, Minbo Su, Yaqi Ding, Xia Chen, Zhenyun Du, Can Jin, Chunfan Huang, Yi Zang, Jia Li, Yechun Xu, Xinwen Chen, Bo Zhang, Kan Ding

## Abstract

Severe acute respiratory syndrome coronavirus 2 (SARS-CoV-2) is the etiological agent responsible for the worldwide coronavirus disease 2019 (COVID-19) outbreak. Investigation has confirmed that polysaccharide heparan sulfate can bind to the spike protein and block SARS-CoV-2 infection. Theoretically, similar structure of nature polysaccharides may also have the impact on the virus. Indeed, some marine polysaccharide has been reported to inhibit SARS-Cov-2 infection *in vitro*, however the convinced targets and mechanism are still vague. By high throughput screening to target 3CLpro enzyme, a key enzyme that plays a pivotal role in the viral replication and transcription using nature polysaccharides library, we discover the mixture polysaccharide 375 from seaweed *Ecklonia kurome* Okam completely block 3Clpro enzymatic activity (IC_50_, 0.48 µM). Further, the homogeneous polysaccharide 37502 from the 375 may bind to 3CLpro molecule well (kD value : 4.23 × 10^−6^). Very interestingly, 37502 also can potently disturb spike protein binding to ACE2 receptor (EC_50_, 2.01 µM). Importantly, polysaccharide 375 shows good anti-SARS-CoV-2 infection activity in cell culture with EC_50_ values of 27 nM (99.9% inhibiting rate at the concentration of 20 µg/mL), low toxicity (LD_50_: 136 mg/Kg on mice). By DEAE ion-exchange chromatography, 37501, 37502 and 37503 polysaccharides are purified from native 375. Bioactivity test show that 37501 and 37503 may impede SARS-Cov-2 infection and virus replication, however their individual impact on the virus is significantly less that of 375. Surprisingly, polysaccharide 37502 has no inhibition effect on SARS-Cov-2. The structure study based on monosaccharide composition, methylation, NMR spectrum analysis suggest that 375 contains guluronic acid, mannuronic acid, mannose, rhamnose, glucouronic acid, galacturonic acid, glucose, galactose, xylose and fucose with ratio of 1.86 : 9.56 : 6.81 : 1.69 : 1.00 : 1.75 : 1.19 : 11.06 : 4.31 : 23.06. However, polysaccharide 37502 is an aginate which composed of mannuronic acid (89.3 %) and guluronic acid (10.7 %), with the molecular weight (*Mw*) of 27.9 kDa. These results imply that mixture polysaccharides 375 works better than the individual polysaccharide on SARS-Cov-2 may be the cocktail-like polysaccharide synergistic function through targeting multiple key molecules implicated in the virus infection and replication. The results also suggest that 375 may be a potential drug candidate against SARS-CoV-2.

## Introduction

The Severe Acute Respiratory Syndrome Coronavirus 2 (SARS-CoV-2) has made a pandemic of Coronavirus Disease 2019 (COVID-19) cross the globe. Up to now, this virus has spread more than 200 countries. Around 22 million confirmed infection and 1500,000 have died in the past 6 months and the number is increasing (Dong, Du, & Gardner, 2020). Keeping social distancing is a functional strategy to slow down the infection due to the absence of vaccine or during the coming vaccine injection period and effective medicine. Currently, only one anti-SARS-Cov-2 agent, remdesivir, has been approved by FDA for the treatment of adult COVID-19 patients (Beigel et al., 2020). Scientific research institutions and pharmaceutical companies are trying to understand the mechanism of SARS-CoV-2 infection and potential antivirus drug to treat COVID-19. A chymotrypsin-like cysteine protease called 3C-like protease (3CLpro) and papain-like protease (PLpro) are required to process polyproteins into mature nonstructural proteins such as RNA-dependent RNA polymerase (RdRp) and helicase, which are essential for viral transcription and replication (Su et al., 2020). Shailendra K. Saxena, et al, found that SARS-CoV-2 had only 12.8 % of difference with SARS-CoV in S protein and has 83.9 % similarity in minimal receptor-binding domain with SARS-CoV (Kumar, Maurya, Prasad, Bhatt, & Saxena, 2020). In 2003, it had been identified that angiotensin-converting enzyme 2 (ACE2) could efficiently bind to the S1 domain of the SARS-CoV S protein (Li et al., 2003). The S protein is a heavily glycosylated protein, which possess 22 potential N-glycosylation sites and facilitates attachment, entry and membrane fusion (Shajahan, Supekar, Gleinich, & Azadi, 2020). Previous researchers showed that several viruses interacted with sialic acids located on the ends of glycans in glycolipids and glycoproteins surrounding the cells. Some other viruses might interact with heparan sulfate (HS) that is attached to cell membrane (Milewska et al., 2014) or extracellular matrix proteoglycans (Lindahl, Couchman, Kimata, & Esko, 2015).

Latest research found that SARS-CoV-2 entry the human cell through binding of the virus spike (S) protein to angiotensin-converting enzyme 2 (ACE2) and cellular HS on the surface of the host cell (Clausen et al., 2020). Hence, blocking the viral transcription, replication and interfering the binding of SARS-CoV-2 and human cells targeting the glycan on ACE2 or spike (S) protein are rational strategies to fight SARS-Cov-2 infection. Interestingly, traditional Chinese medicine has been paid more attention for antivirus clinical drug application during SARS-CoV and SARS-CoV-2 spreading (Leung, 2007; H. Luo et al., 2020; Wen et al., 2011). Indeed, Baicalin and baicalein, two ingredients of Chinese traditional patent medicine Shuanghuanglian, were characterized as the first noncovalent, nonpeptidomimetic inhibitors of SARS-COV-2 3CLpro and exhibited potent antiviral activities in a cell-based system (Su et al., 2020) . However, the detailed mechanism underlying active components against the virus is still vague.

Traditional Chinese medicine *Ecklonia kurome*, named Kunbu in China, is a seaweed of the *Laminariaceae*, belonging to the genus *Laminariales* (Kuda, Kunii, Goto, Suzuki, & Yano, 2007). The high pharmaceutical value of this seaweed is at least partially relied on the biomacromolecule polysaccharide. Different from other general polysaccharide, the polysaccharide in *Ecklonia kurome* is mainly present on cell wall of brown algae plants and has a sulfate group at the end of its molecular chain, which may make it have significance bio-activities (Li et al., 2020). For instance, a modified sulfate polysaccharide extracted from *Ecklonia kurome* has anti-angiogenesis anti-tumor effect (Kuda et al., 2007). Alginate, a mainly acid polysaccharide in *Ecklonia kurome*, is a liner anionic polymer of β-(1-4)-D-mannuronic acid and of its C-5 epimer, α-(1-4)-L-guluronic acid. They consist of alternate homopolymeric blocks of poly-β-(1-4)-D-mannuronic acid (M), and poly-α-(1-4)-L-guluronic acid (G), and of heteropolymeric blocks with random arrangements of both monomers (Fujihara & Nagumo, 1993). Previous research revealed that alginate had the effect on anti-tumor and immunoenhancement (de Sousa et al., 2007; Luo, Wu, Lou, Zhao, & Yang, 2020), however no anti-virus effect was reported.

In this study, we firstly extracted crude polysaccharide, named 375 from *Ecklonia kurome* and examine the bioactivity against SARS-Cov-2 virus. Then we isolated and purified crude polysaccharide and further characterized the structure of one homogeneous polysaccharide 37502 from *Ecklonia kurome*. The molecular weight (*Mw*), monosaccharide composition, infrared spectroscopy (IR) and nuclear magnetic resonance (NMR) spectra were employed to analyze the structure. Eventually, we further examined anti-SARS-CoV-2 bioactivity by targeting 3CL pro, S1 and ACE2 molecules.

## Results and discussion

### Isolation, purification and monosaccharide composition analysis of 37502

The crude polysaccharide, 375 (200 mg) was fractioned by DEAE Sepharose™ Fast Flow. 37501 (20 mg, yield: 10 %), 37502 (44 mg, yield: 22 %) and 37503 (35 mg, yield: 17 %) were obtained by elution with distilled water, 0.2 M NaCl and 0.3 M NaCl (**Fig. 1**). 375 content 65 % neutral polysaccharide, 5 % protein and 28 % uronic acid. 37501 content 12 % neutral polysaccharide and 7.7 % uronic acid. 37502 content 16.4 % neutral polysaccharide and 32.0 % uronic acid. The sugar content, protein and uronic acid of 37503 were 40.3 %, 2.3 % and 11.9 %, respectively. The homogeneity of 37502 was determined by high performance gel permeation chromatography (HPGPC) that showed a single symmetrical peak with the *Mw* of 27.9 kDa (**Fig. S1A, Supplementary data**). The monosaccharide compositions of 375, 37501, 37502 and 37503 were analyzed by HPLC after PMP derivatization with PMP in parallel with monosaccharides standards. Compared with standards, the monosaccharide composition of 375, 37501 and 37503 were shown in **Fig. S2**, the crude polysaccharide 375 contained guluronic acid (3.0 %), mannuronic acid (15.3 %), mannose (10.9 %), rhamnose (2.7 %), glucouronic acid (1.6 %), galacturonic acid (2.8 %), glucose (1.9 %), galactose (17.7 %), xylose (6.9 %) and fucose (36.9 %). 37501 mainly contained guluronic acid (1.4 %), mannuronic acid (3.6 %), mannose (14.6 %), rhamnose (3.6 %), glucouronic acid (2.0 %), galactose (28.1 %), xylose (7.37 %) and fucose (39.1 %). 37503 was composed of mannose (30.1 %), rhamnose (9.6 %), glucose (3.4 %) galactose (7.3 %), xylose (10.9 %) and fucose (36.9 %). Relatively, the monosaccaride composition of 37502 was more simple, it was composed of mannuronic acid (89.3 %) and guluronic acid (10.7 %) as shown in (**Fig. 2**). More structure information was revealed by FT-IR and NMR spectra.

**Fig. 1.**
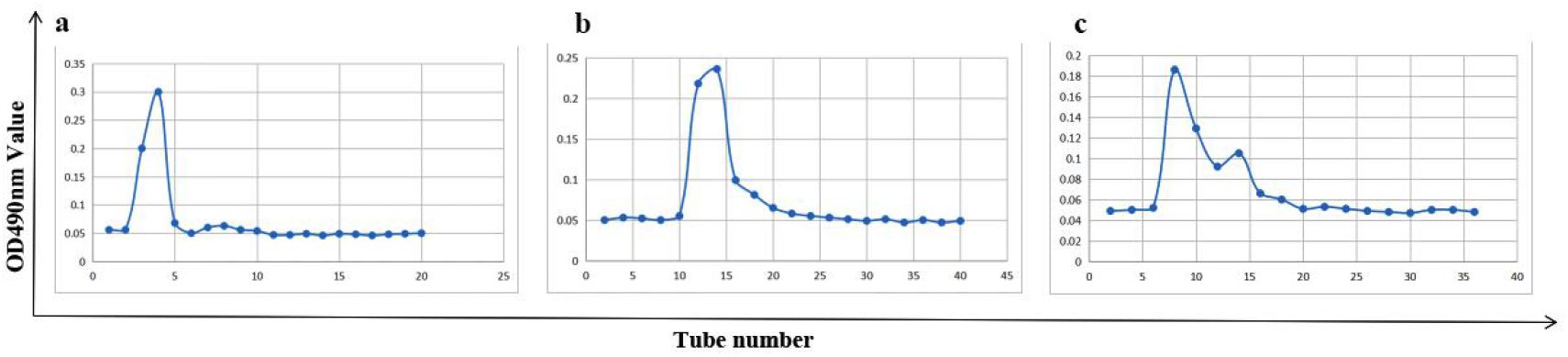
Elution profile of 375 on DEAE Sepharose Fast Flow with different NaCl; a. distilled water elution profile; b. 0.2 M NaCl elution profile; c. 0.3 M NaCl elution profile. concentration.

**Fig. 2.**
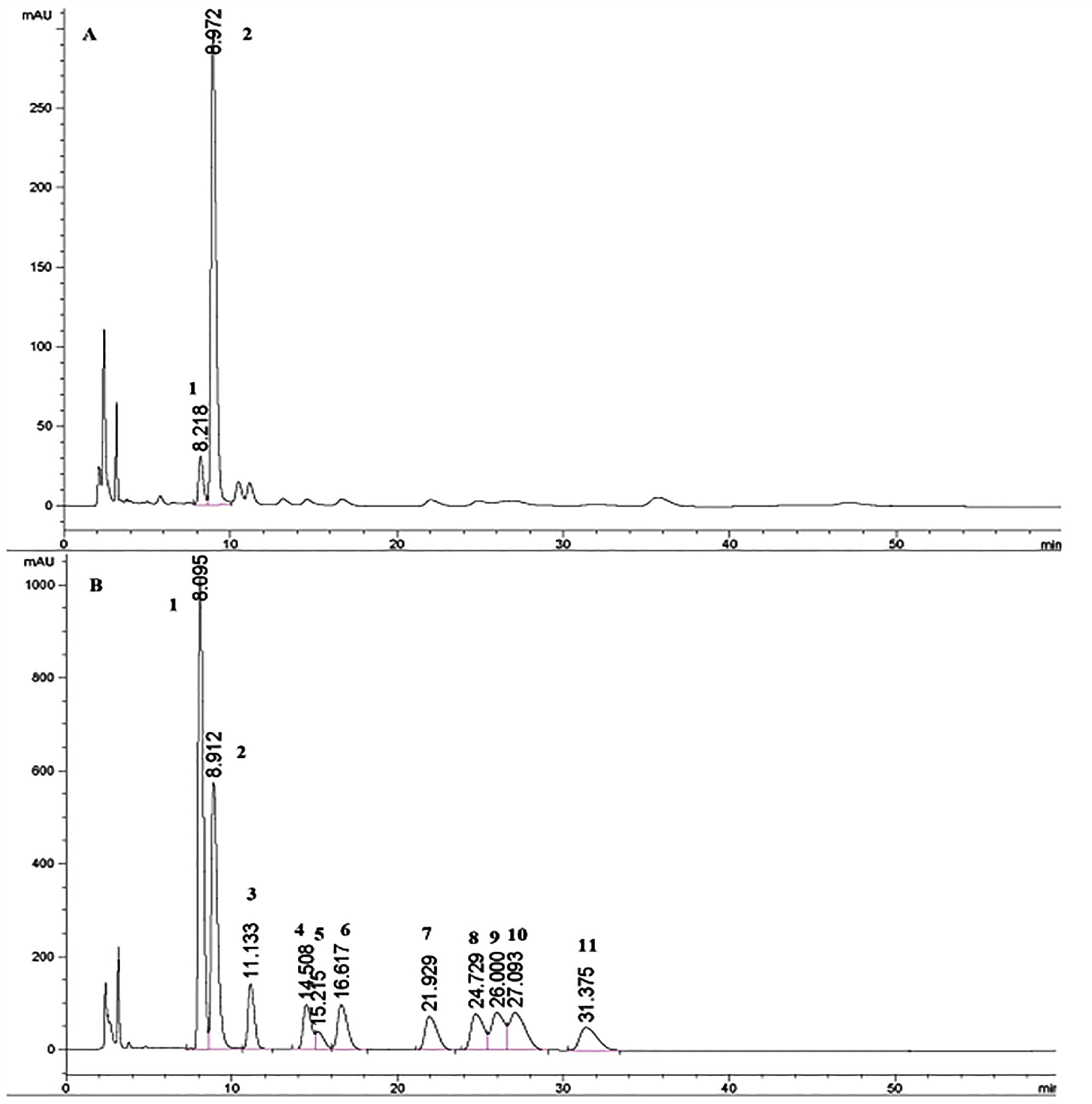
Determination of monosaccharide composition of 37502 by HPLC. A. 37502; B. Monosaccharide standards (1. guluronic acid; 2. mannuronic acid; 3. Mannose; 4. Rhamnose; 5. Glucouronic acid; 6. Glactouronic acid; 7. Glucose; 8. Galactose; 9. Xylose; 10. Arabinose; 11. Fucose).

### FT-IR spectrum

FT-IR spectrum of 37502 showed the typical absorption bands of the polysaccharide (**Fig. 3**). 3453.88 cm^-1^ was assigned to the O-H stretching bands. 2935.13 cm^-1^ was from the stretching bands of C-H group. 1616.06 cm^-1^ and 1417.42 cm^-1^ could been reassigned to the asymmetrical and symmetrical stretching vibration of COO-, respectively, which confirmed the presence of uronic acid. The absorption at 1035.59 cm^-1^ showed the existence of the stretching vibration of C-C. 956.52 cm^-1^ was the stretching vibration of uronic acid residues. 890.95 cm^-1^ was the variable angular vibration of C1-H of β-mannuronic acid (Mao, Li, Gu, Fang, & Xing, 2004). 821.53 cm^-1^ was the special absorption peak of mannuronic acid and 792.12 cm^-1^ was the special absorption peak of guluronic acid (Lin et al., 2007).

**Fig. 3.**
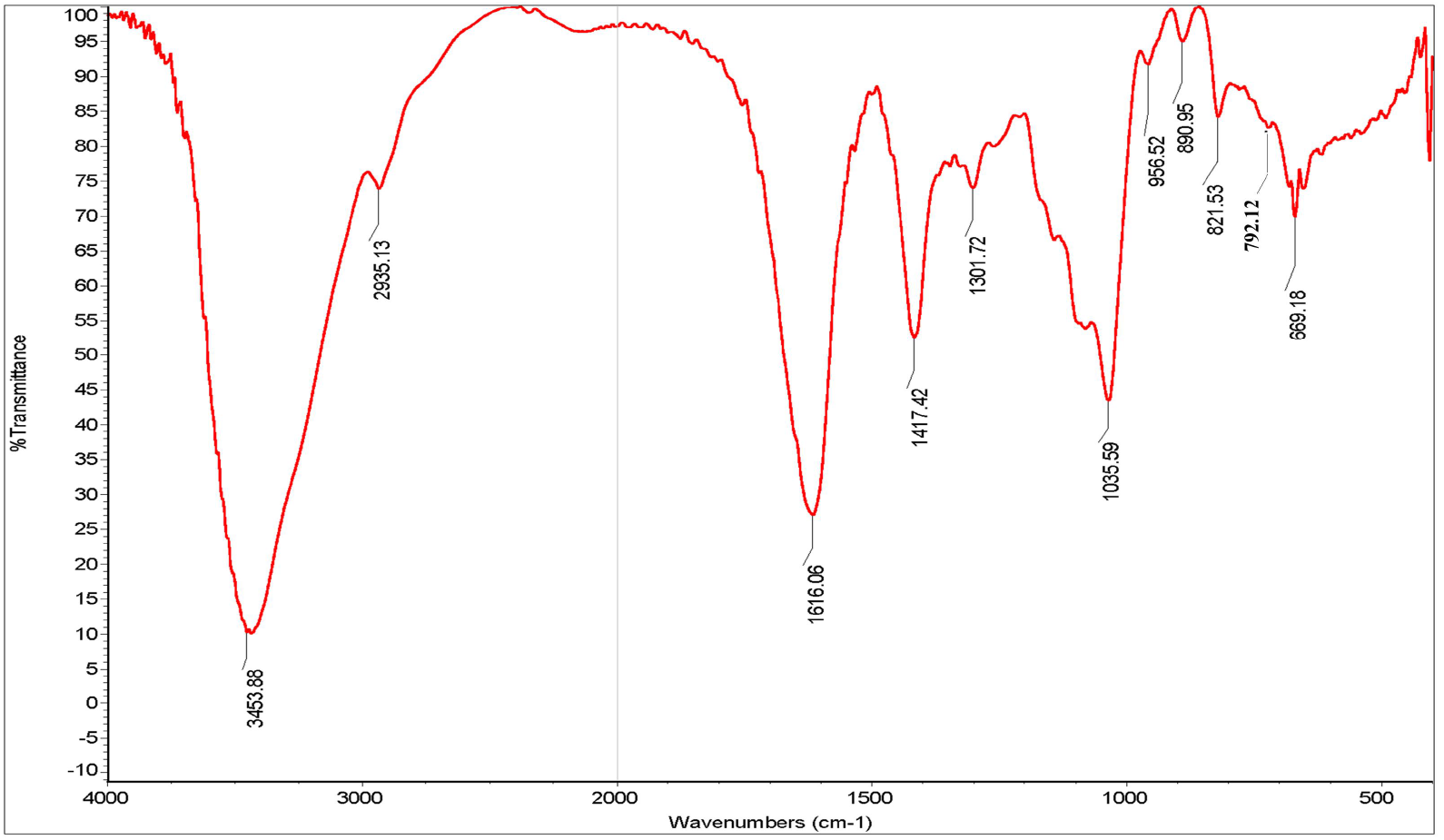
FT-IR spectrum of 37502

### Linkage pattern analysis

In order to determine the linkage types, the methylation method was employed. First, 37502 was reduced with 1-Cyclo-hexyl-3-(2-mopholinoethyl) carbodiimidemetho-p-toluenesulfonate (CMC) and the reduced production was further methylated. Result revealed that there are mainly 2,3,6-Me_3_-Man and little 2, 3, 4, 6-Me_4_-Man, indicating that 37502 was a linear 1, 4-linked mannosan. More structure information will be shown by NMR spectra.

### NMR spectral analysis

^13^C NMR spectra of 375, 37502 and 37503 were compared in **Fig. S3** and the ^1^H, ^13^C NMR spectra of 37502 were shown in **Fig. 4**. In the ^13^C NMR spectra of 37502 (**Fig. 4A**), the signals at δ 102.45 ppm and 101.22 ppm were assigned to C1 of 1, 4-linked α-L-guluronic acid and 1, 4-linked β-D-mannuronic acid, respectively (Heyraud et al., 1996). The strong signals at δ 65.87, δ 78.46 and δ 68.57 were ascribed to C-2, C-3 and C-5 of the 1,4-linked α-L-guluronic acid. The signals at δ 71.12, δ 72.55 and δ 76.69 were ascribed to C-2, C-3 and C-5 of the 1, 4-linked β-D-mannuronic acid. The signals at δ 81.2 and δ 79.06 were assigned to C-4 of 1,4-linked α-L-guluronic acid and 1, 4-linked β-D-mannuronic acid, respectively (Schürks, Wingender, Flemming, & Mayer, 2002).

**Fig. 4.**
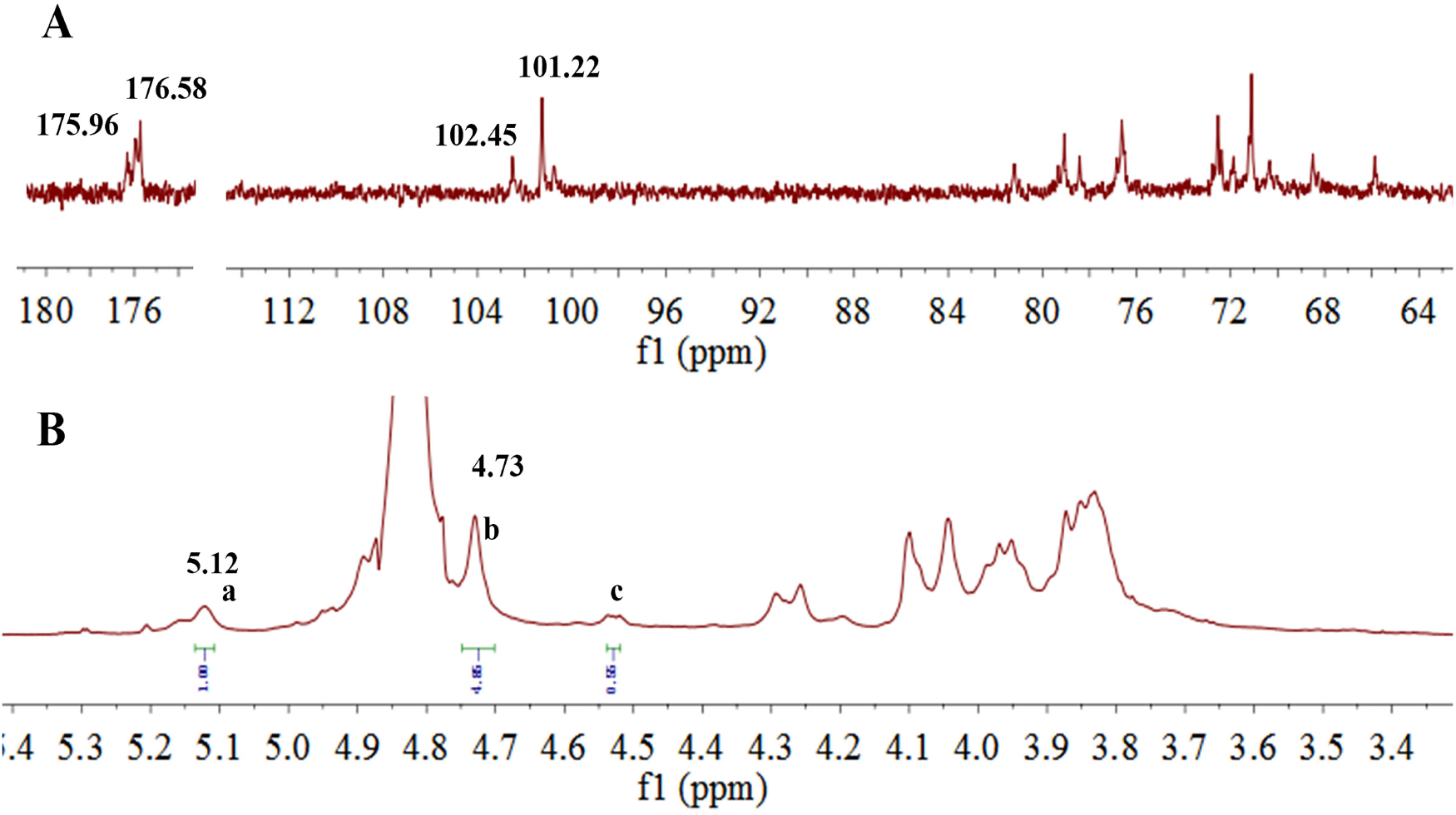
^1^H NMR spectrum (A) and ^13^C NMR spectrum (B) of 37502.

In the ^1^H NMR (**Fig. 4B**) and HSQC (**Fig. 5B**) spectra, δ 5.12 was assigned to H-1 of 1,4-linked α-L-guluronic acid, which correlated to C-1 of 1,4-linked α-L-guluronic acid (102.45 ppm). δ 4.73 was assigned to H-1 of 1, 4-linked β-D-mannuronic acid correlated to C-1 of 1, 4-linked β-D-mannuronic acid (101.22 ppm). In ^1^H-^1^H COSY (**Fig. 5A**) spectra, δ 4.04 was assigned to H-2 of 1,4-linked α-L-guluronic acid, which correlated to H-1 (δ 5.12) of 1,4-linked α-L-guluronic acid. δ 4.10 was assigned to H-2 of 1, 4-linked β-D-mannuronic acid, which correlated to H-1 (δ 4.73) of 1, 4-linked β-D-mannuronic acid. Other signals were listed in **Table.1**.

**Table.1.** ^1^H NMR and ^13^C NMR chemical shifts (ppm) assignments for 37502.

| Residues |  | 1 | 2 | 3 | 4 | 5 | 6 |
| --- | --- | --- | --- | --- | --- | --- | --- |
| 1, 4-linked guluronic acid (G) | C | 102.45 | 65.87 | 78.46 | 81.2 | 68.57 | 176.58 |
|  | H | 5.12 | 4.04 | 3.95 | 4.29 | 4.53 |  |
| 1, 4-linked mannuronic acid (M) | C | 101.22 | 71.12 | 72.55 | 79.06 | 76.69 | 175.96 |
|  | H | 4.73 | 4.10 | 3.83 | 3.93 | 3.87 |  |

**Fig. 5.**
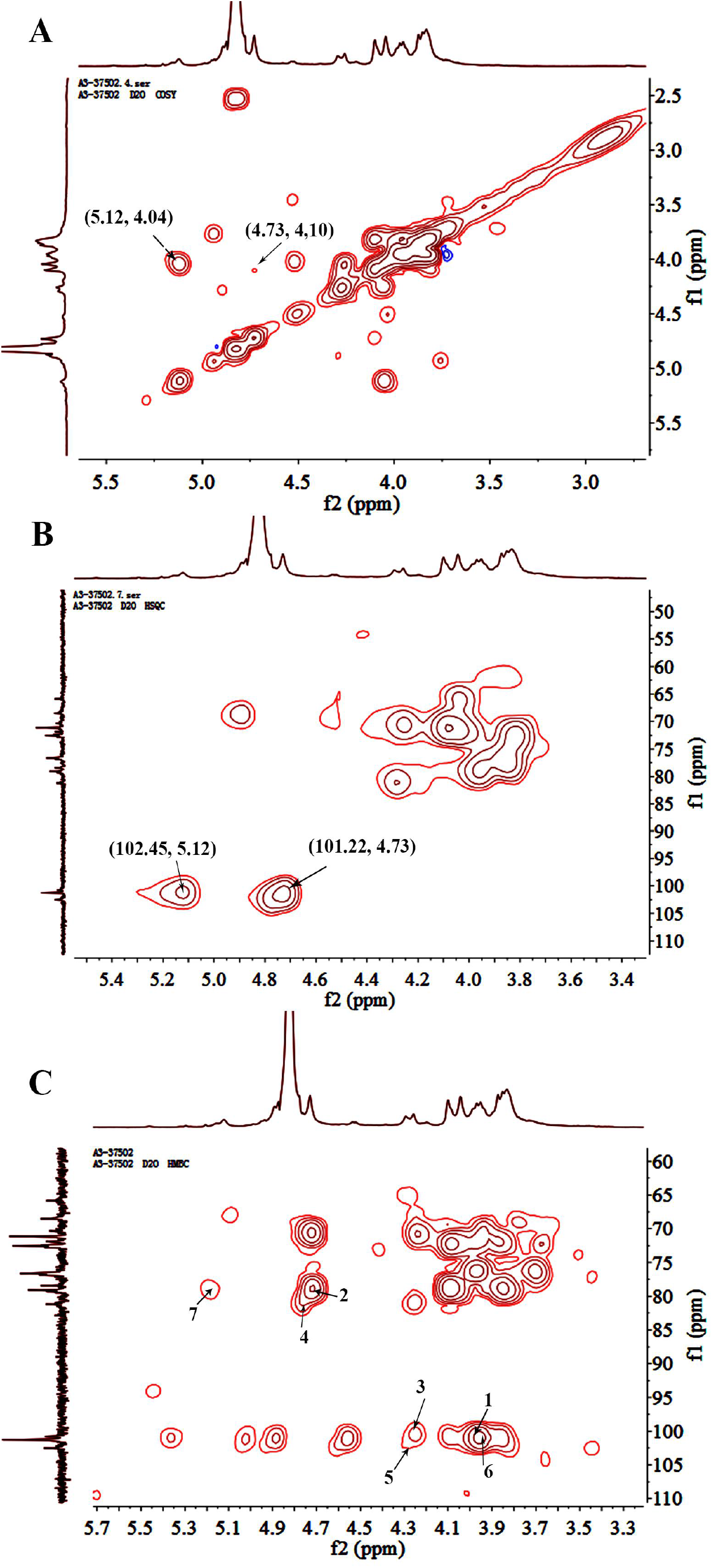
Two dimensional spectra of 37502; A. 1H-1H COSY spectrum; B. HSQC spectrum; C. HMBC of spectrum.

HMBC spectrum was employed to analyze the structure backbone and the substitution sites. In **Fig. 5C**, the strong peak 1 (101.22/3.93) represented the correlation between C-1 and H-4 of 1, 4-linked mannuronic acid. The cross peak 2 (79.06/4.73) represented the correlation between C-4 and H-1 of 1, 4-linked mannuronic acid. These two peaks showed the existence of liner 1, 4-linked mannuronic acid. The cross peak 3 (101.22/4.29) showed the correlation between C-1 of 1, 4-linked mannuronic acid and H-4 of the neighboring 1, 4-linked guluronic acid. The peak 4 (81.2/4.73) represent the correlation between C-4 of 1, 4-linked guluronic acid and H-1 of the neighboring 1, 4-linked mannuronic acid. These results suggested that guluronic acid may linked to the liner mannuronic acid. The cross peak 5 (102.45/4.29) showed the correlation between C-1 of 1, 4-linked guluronic acid and H-4 of neighboring 1, 4-linked guluronic acid, which showed the existence of liner 1, 4-linked guluronic acid. The cross peak 6 (102.45/3.93) overlapped by peak 1 showed the correlation between C-1 of 1, 4-linked guluronic acid and H-4 of neighboring 1, 4-linked mannuronic acid. The cross peak 7 (79.06/5.12) also showed the correlation between C-4 of 1, 4-linked mannuronic acid and H-1 of 1, 4-linked guluronic acid. It showed the existence of heteropolymeric blocks with random arrangements of both mannuronic acid and guluronic acid.

### Putative structure of 37502

Based on the above results, we proposed that 37502 is an alginate consisting of different dimer structures (MM, MG, GM and GG). The molar ratio (F) of dimer structures was speculated according to the integral area (*I*) of **a, b, c** in ^1^H-NMR and following formula: F_G_ = *I***a**/(*I***b**+*I***c**), F_GG_ = *I***c**/(*I***b**+*I***c**), F_GG_ +F_GM_ = F_G_, F_MM_ + F_MG_ = F_M_, F_MG_ = F_GM_. In ^1^H-NMR of 37502, the integral area of **a, b, c** was 1.0, 4.85 and 0.55, respectively. So, F_G_ = 0.19; F_M_ = 0.9; F_MM_ = 0.81; F_GG_ = 0.1; F_GM_ = F_GM_ = 0.09. The molar ratio (0.19: 0.9) of between F_G_ and F_M_ was consisted of the results of monosaccharide composition. The ratio of F_MM_ was highest and the ratio of F_GG_, F_MG_ and F_GM_ was similar. Finally, the possible structure of 37502 (*Mw*: 27.9 kDa) was shown in **Fig. 6**.

**Fig. 6.**
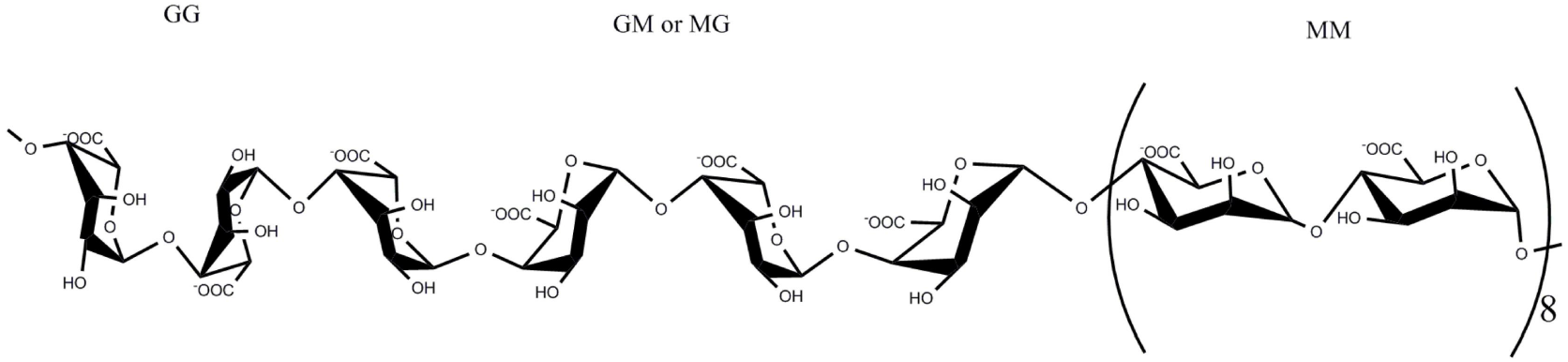
Proposed structure of 37502

### Crude polysaccharide 375 was a potent inhibitor of SARS-COV-2 3CLpro

People showed that marine polysaccharide may inhibit SARS-Cov-2 infection (Song S, et al. 2020), however the convinced targeting molecule and mechanism are still unknown. SARS-CoV-2 includes two open reading frames ORF1a and ORF1b (Yin et al., 2020). ORF1a encodes two cysteine proteases, a papain-like protease (PLpro) and a 3 C-like protease (3CLpro). Scientists have provided evidences that main protease (3CLpro) is one of the good targets to discover new antiviral agents before vaccines are available (Derosa, Maffioli, D’Angelo, & Di Pierro, 2020). To explore more potent leading compound against SARS-Cov-2 by targeting key molecule in RNA synthesis of the virus, recombinant SARS-CoV-2 3CLpro was firstly expressed and purified from *Escherichia coli* (Xue et al., 2007; Yang et al., 2003). A fluorescently labeled substrate MCA-AVLQSGFR-Lys (Dnp) -Lys-NH2, derived from the N-terminal autocleavage sequence from the viral protease was designed and synthesized for the enzymatic assay. Then the binding test targeting 3CLpro was examined. The results showed that 375 might potently inhibit SARS-CoV-2 3CLpro activity (Fig. 7). Further we used a fluorescence resonance energy transfer (FRET) based cleavage assay to determine the median inhibitory concentration (IC_50_) values. The results also revealed good inhibitory potency, with IC _50_ values of 0.48 ± 0.1 µM (Fig. 7). It implies that 375 may contain the bioactive components to inhibit SARS-CoV-2 replication and infection. Based on fact that 375 is a crude polysaccharide, the results inspire us to explore which bioactive components from crude polysaccharide 375 may contribute the effect against SARS-Cov-2 and their underlying mechanism.

**Fig. 7.**
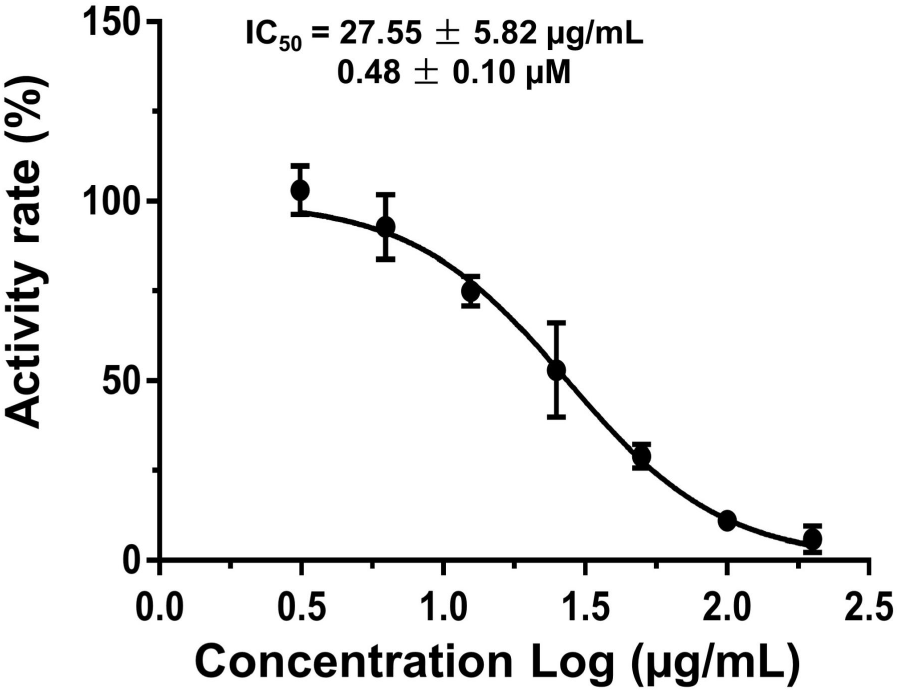
Inhibitory activity profiles of 375 against SARS-CoV-2 3CLpro

### 37502 may bind to SARS-CoV-2 3CLpro and disturb the interaction between SARS-CoV-2-S1 and ACE2

To explore which compound from 375 may interfere with 3CLpro enzyme activity, isothermal titration calorimetry (ITC) method was employed (Su et al., 2020). Result showed that. 37502, one homogeneous polysaccharide from 375 could bind to SARS-CoV-2 3CLpro protein very well (Fig. 8A). The Kd value was 4.23 × 10^−6^ M (**Fig. 8A**). This result suggested that 375 or 37502 might interfere with the replication of SARS-CoV-2 in some way.

**Fig. 8.**
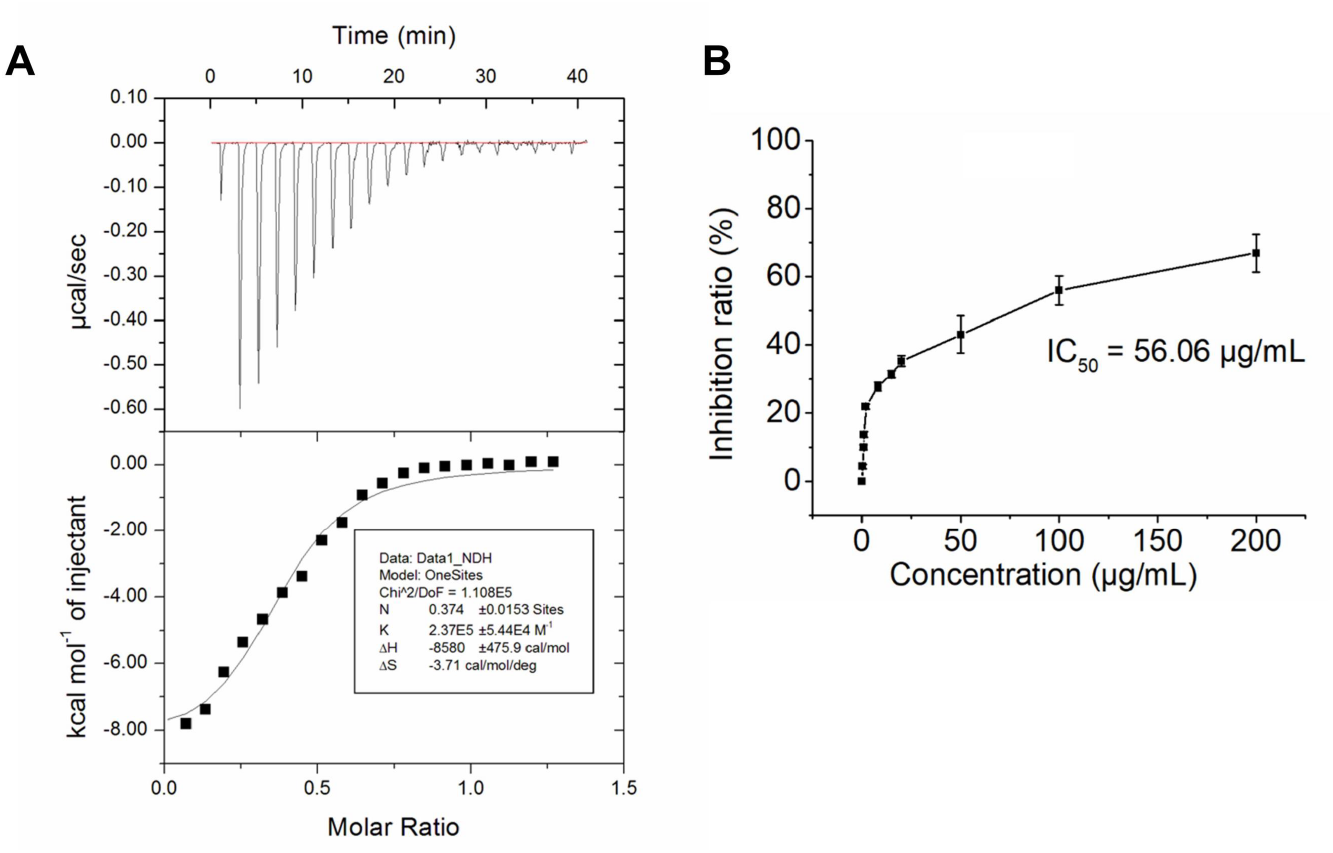
(A) Binding test of polysaccharide 37502 with SARS-CoV-2 3CLpro by ITC. (B) Competitive intervention of polysaccharide 37502 on S1 protein and ACE2 by ELISA.

The spike protein of SARS-CoV-2 shows more than 90% amino acid similarity to that of pangolin and bat CoVs which also use human angiotensin-convert enzyme 2 (ACE2) as a receptor for the virus infection. Receptor binding domain of S protein of SARS-CoV-2 which are processed into two subunits including S1 and S2, can bind with ACE2 as a receptor to invade target cells (Lan et al., 2020). Thus, S protein is very vital for viral invasion. Interestingly, the N-terminal domain of S1 protein has the glycan binding site (Kirchdoerfer et al., 2016). This implies S protein can bind with carbohydrate. To further understand the bioactivity of 375 against SARS-Cov-2, purified homogeneous polysaccharide 37502 from 375 was employed to examine whether it might disturb the binding between S1 protein and ACE2 by ELISA method. The results showed that polysaccharide 37502 could effectively impede the binding of S1 protein with ACE2. The median inhibitory concentration (IC_50_) was 56.06 µg/mL (**Fig. 8B**). This result suggested that 37502 might had the potential to block SARS-Cov-2 infection through disturbing the S protein binding to ACE2 receptor.

### 375 and 37503 exhibit anti-viral effect on SARS-CoV-2

The above results inspired us to explore whether those polysaccharides might really block SARS-Cov-2 replication. Firstly, the inhibition effect of native polysaccharide 375 was examined. Surprisingly, as shown in **Fig. 9A**, polysaccharides 375 potently inhibit SARS-Cov-2 *in vitro* with a EC_50_ value about 27 nM (or 1.56 µg/mL) (**Fig. 9A**). However, the toxicity of 375 on mice is low and the LD_50_ is 136 mg/kg (**Table S1, Supplementary Data**). To further understand which component from native polysaccharide 375 is contributing to inhibit effect on SARS-Cov-2, quantitative real-time polymerase chain reaction (qRT-PCR) was also employed to monitor the antiviral activity of the purified polysaccharide 37501, 37502 and 37503 from 375. The result showed that polysaccharide 37501 and 37503 exhibited antivirus effect on SARS-CoV-2 (**Fig. 9B**). Although 37502 might attenuate 3CLpro enzymatic activity significantly (**Fig. 8A**), it had no significant direct effect against the active virus replication (**Fig. 9B**). To examine whether purified polysaccharide 37503 have stronger bioactivity than that of native polysaccharide 375, EC_50_ of 37503 against SARS-Cov-2 was also measured. Interestingly, effect of 37503 is significant feeble than that of 375, while EC_50_ of 37503 is 0.89 µM (or 11.07 µg/mL) (**Fig. 9C**). This indicated the effects of 375 against SARS-Cov-2 were cocktail-like synergistic contribution of combined components of 37501, 37502, and 37503, the impact of individual homogeneous polysaccharide might be weaker although. In brief, although polysaccharide 375 and 37503 are probably the potential drug candidate for inhibiting the SARS-CoV-2 infection, obviously cocktail-liked crude polysaccharide 375 is the best option for anti-SARS-Cov-2 new drug development. Following, we will focus on 375 and 37503 for further investigation.

**Fig. 9.**
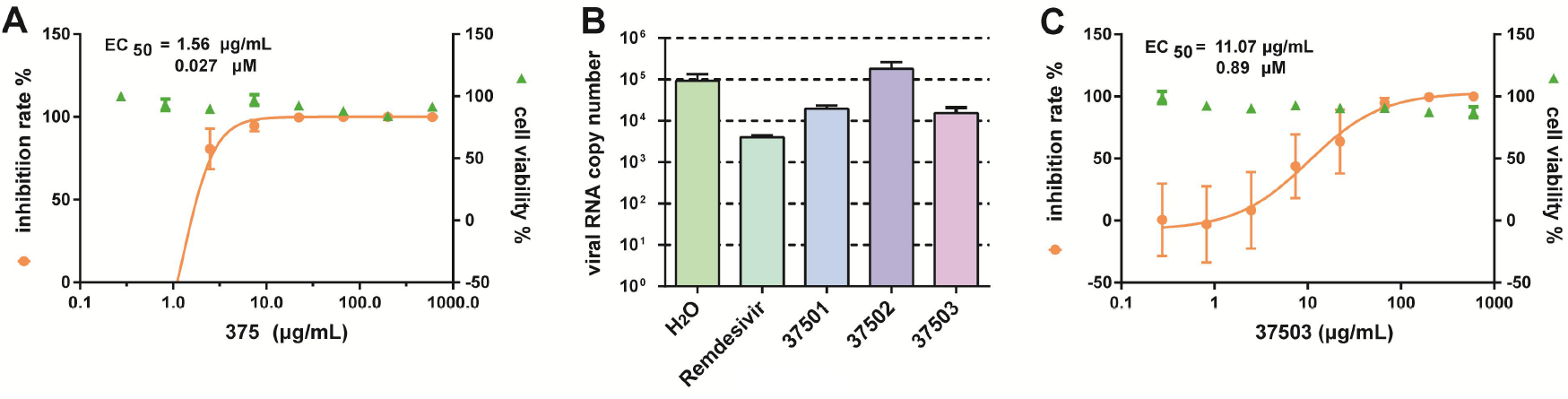
*In vitro* inhibition of polysaccharides against SARS-CoV-2. (A) EC_50_ of crude polysaccharide 375 against SARS-CoV-2; (B) viral RNA copy number was detected by qPCR after the treatment of solvent (H_2_O) control, Remdesivir positive control, crude polysaccharide 375 and their fragmentation37501, 37502 and 37503. (C) EC_50_ of homogenous polysaccharide 37503 against of SARS-CoV-2.

## Experimental

### Materials and reagents

DEAE Sepharose Fast Flow was obtained from GE healthcare. Dimethyl sulfoxide (DMSO) was from E. Merck. U.S.A. Sodium borohydride (NaBH_4_) and iodomethane were obtained from Sinopharm Chemical Reagent Co. Ltd. Standard monosaccharides were purchased from Shanghai Aladdin Bio-Chem Technology Co. Ltd. Other reagents were analytical grade and from Sinopharm Chemical Reagent Co. Ltd. (Shanghai, China).

### Determination of physicochemical property of polysaccharides

The carbohydrate content was determined by the PhOH-H_2_SO_4_ method using glucose as a standard (Dubios, Gilles, Hamilton, Rebers, & Smith, 1956). Protein content was evaluated using a BCA protein assay kit (Beyotime Biotechnology, China). Uronic acid content was determined by the meta-hydroxydiphenyl method using galacturonic acid as a standard. All the measurements were repeated three times. Novostar microplate reader was employed to detect the absorbance at OD_490_ for sugar, OD_520_ for uronic acid and OD_562_ for protein.

### Homogeneous polysaccharide preparation

Crude polysaccharide was fractionated on a DEAE Sepharose Fast Flow column. 200 mg polysaccharide was dissolved in 20 mL distilled water and centrifuged for each time. The supernatant was applied to a DEAE Sepharose Fast Flow column and eluted with distilled water, 0.05, 0.1, 0.2 and 0.3 M NaCl solutions stepwise. The solution was pooled according to the elution profile based on phenol-sulfuric acid method. Then the fraction eluted with distilled water, 0.2 M NaCl and 0.3 M NaCl were collected, followed by concentration, dialysis with distill water and freeze-dried.

### Homogeneity and molecular weight

The homogeneity and molecular weight (*Mw*) were examined by HPGPC (high performance gel permeation chromatography) method on Agilent 1260 HPLC (Santa Clara, CA, USA) system equipped with two series-connected columns (UItrahydrogel™ 2000 and 500). The columns were calibrated by pullulans standards. 0.1 mol/L NaNO_3_ was used as an eluent and the flow rate was maintained at 0.6 mL/min. The column temperature was maintained at 40.0 °C ± 0.1 °C. The samples were prepared with mobile phase as 0.2 % (W/V) solution. 20 µL of sample was injected for analysis.

### Monosaccharide composition analysis

The method of monosaccharide composition was PMP pre-column derivatisation based on the previous reported (Dai et al., 2010).

### FT-IR spectrum and NMR analysis

The IR spectra was determined according to the previous report (Cong, Xiao, Liao, Dong, & Ding, 2014). 2 mg native polysaccharide was mixed with dried KBr powder and pressed into pellet, then scanned from 4,000 to 600 cm^-1^ for the analysis.

For the NMR analysis, 30 mg of the sample was deuterium-exchanged and dissolved in 0.5 mL D_2_O. The ^1^HNMR, ^13^CNMR, ^1^H-^1^H COSY, HSQC and HMBC were measured at 25 °C with acetone as internal standard (δ_C_ = 31.5, δ_H_ = 2.29). NMR spectra were recorded on a Bruker AVANCE III NMR spectrometer operating at 500 MHz.

### Methylation analysis

10 mg sample was methylated with a modified method from Ciucanu and Kerek. Briefly, the sample was dried overnight with P_2_O_5_ and dissolved in DMSO (2 mL) and 100 mg powdered NaOH was added into the reaction bulb and stirred at room temperature for 3 h. Iodomethane (0.5 mL) was added dropwise within 45 min in ice-cold water bath. The mixture was stirred for 2.5 h at room temperature in a dark place and then 1 ml deionized water was added to terminate the reaction. The redundant CH_3_I was removed by evaporation under depressed pressure. The solution was extracted by 15 ml CHCl_3_ and 15 ml water (1:1, v/v) and the organic phase was washed with deionized water for three times. After removing the residual water by adding Na_2_SO_4_, the organic phase was concentrated to get the methylated polysaccharide. Then the product was hydrolyzed with 2 M TFA for 2 h at 110 °C and converted into the partially methylated alditol acetates (PMAA) and analyzed by GC-MS. The GC-MS program designed for methylation analysis was based on the reported method (Cong et al., 2014).

### Uronic acid reduction

The method was based on the reported method (Taylor & Conrad, 1972). In brief, 40 mg polysaccharide was dissolved in 40 mL H_2_O. CMC (600 mg) was added and pH was kept at 4.75 with 0.01 M HCl for 2 h. Then 2 M fresh aqueous sodium borohydride (15 mL) was added slowly to the mixture in 45 min and 4 M HCl was added concurrently to keep pH at 7.0. The mixture was stirred for 2 h and dialyzed (1,000 mL × 4) for 24 h at room temperature. Then the retentate was lyophilized to achieve carboxyl reduced polysaccharide, followed by monosaccharide composition and linkage pattern analyses.

### Enzymatic activity and inhibition assays

The enzyme activity and inhibition assays have been described previously (Xue et al., 2007; Yang et al., 2005) . The recombinant SARS-CoV-2 3CLpro (30 nM at a final concentration) was mixed with serial dilutions of each compound in 80 µL assay buffer (50 mM Tris-HCl, pH 7.3, 1 mM EDTA) and incubated for 10 min. The reaction was initiated by adding 40 µL fluorogenic substrate with a final concentration of 20 µM. After that, the fluorescence signal at 320 nm (excitation)/405 nm (emission) was immediately measured every 30 s for 10 min with a Bio-Tek Synergy4 plate reader. The V_max_ of reactions added with compounds at various concentrations compared to the reaction added with DMSO were calculated and used to generate IC_50_ curves. For polysaccharide 375, IC_50_ values against SARS-CoV-2 3CLpro were measured at 7 concentrations and three independent experiments were performed. All experimental data was analyzed using GraphPad Prism software.

### ELISA

10 µg/mL ACE2 coating buffer were used to treat the 96 well plate at 4 °C overnight following with 200 µL washing buffer for three times. Then the 96-well plate was blocked by 2% BSA at room temperature for 2 h. After that, 100 µL biotinylated S1 protein was added and incubated at room temperature. At the same time, the positive control and the negative control were set. After incubation for 1 h, the plate was washing for three times and each time for 5 min. Following, 100 µL Streptavidin-HRP was added to final concentration of 200 ng/mL at room temperature and incubated for 1 h. After 1 h incubation, the plate was washed for three times. Then, 100 µL TMB were added and incubated in the dark for 35 min. Finally, 50 µL stop solution were added to stop the reaction and detected the A450 value number by microplate reader (BioTek).

### Isothermal Titration Calorimetry (ITC)

ITC experiment was performed as previously publication (Su et al., 2018). It was conducted with an iTC200 instrument (Microcal, GE Healthcare) at 25 °C, and the resulting data were processed by the supplied MicroCal Origin software package. The concentration of 3CLpro protein used was 600 µM and polysaccharide 37502 used was 300 µM. Titrations were run in triplicate to ensure reproducibility. In all the cases, a single binding site mode was employed and a nonlinear least-squares algorithm was used to obtain best-fit values of the stoichiometry (n), change in enthalpy (*ΔH*), and binding constant (*K*). Thermodynamic parameters were subsequently calculated with the formula ΔG = *ΔH* − *T ΔS* = − *RT* ln *K*, where T, R, Δ G, and *ΔS* are the experimental temperature, the gas constant, the changes in free energy, and entropy of binding, respectively.

### Antiviral test *in vitro*

The experiments related to SARS-CoV-2 are completed at biosafety level 3 (BSL-3) laboratory in the Center for Biosafety Mega-Science, Wuhan, Chinese Academy of Sciences.

SARS-CoV-2 (WIV04) was passaged in Vero E6 cells and tittered by plaque assay. Vero E6 cells were treated with drugs at indicated concentration and infected by SARS-CoV-2 virus at MOI 0.01. After 24 h incubation at 37 °C, supernatants were collected and the viral RNAs were extracted by Magnetic Beads Virus RNA Extraction Kit (Shanghai Finegene Biotech, FG438), and quantified by real-time RT-PCR with Taqman probe targeting to the RBD region of S gene.

## Acknowledgment

This work was supported by Key New Drug Creation and Manufacturing Program (Grant number 2019ZX09735001), National Key R&D Program of China, Ministry of Science and Technology (Grant number 2019YFC1711-000). COVID-19 Emergency Research Project founded by Zhejiang Unversity (2020XGZX080). We are particularly grateful to Tao Du and Lun Wang from the Center for Biosafety Mega-Science for their essential support.

