## supplementary for "Structural characterization of cocktail-like targeting polysaccharides from *Ecklonia kurome* Okam and their anti-SARS-CoV-2 activities *invitro*": supplementary data.docx

**Supplementary information:**

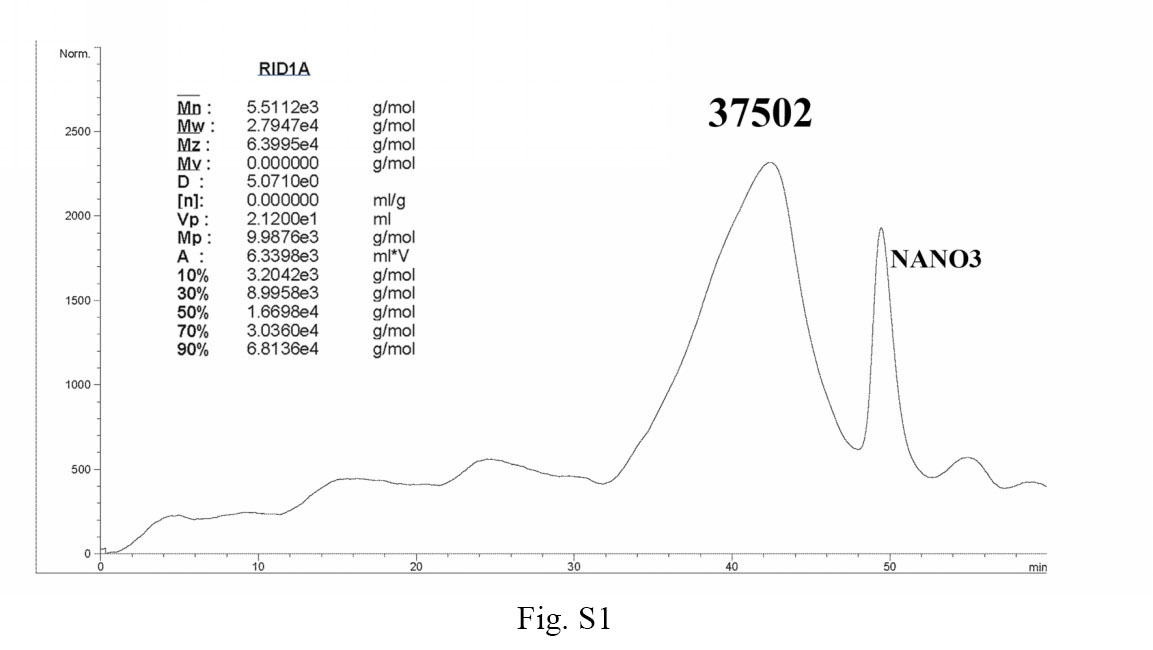

Fig. S1 Homogeneity and molecular weight of polysaccharide 37502

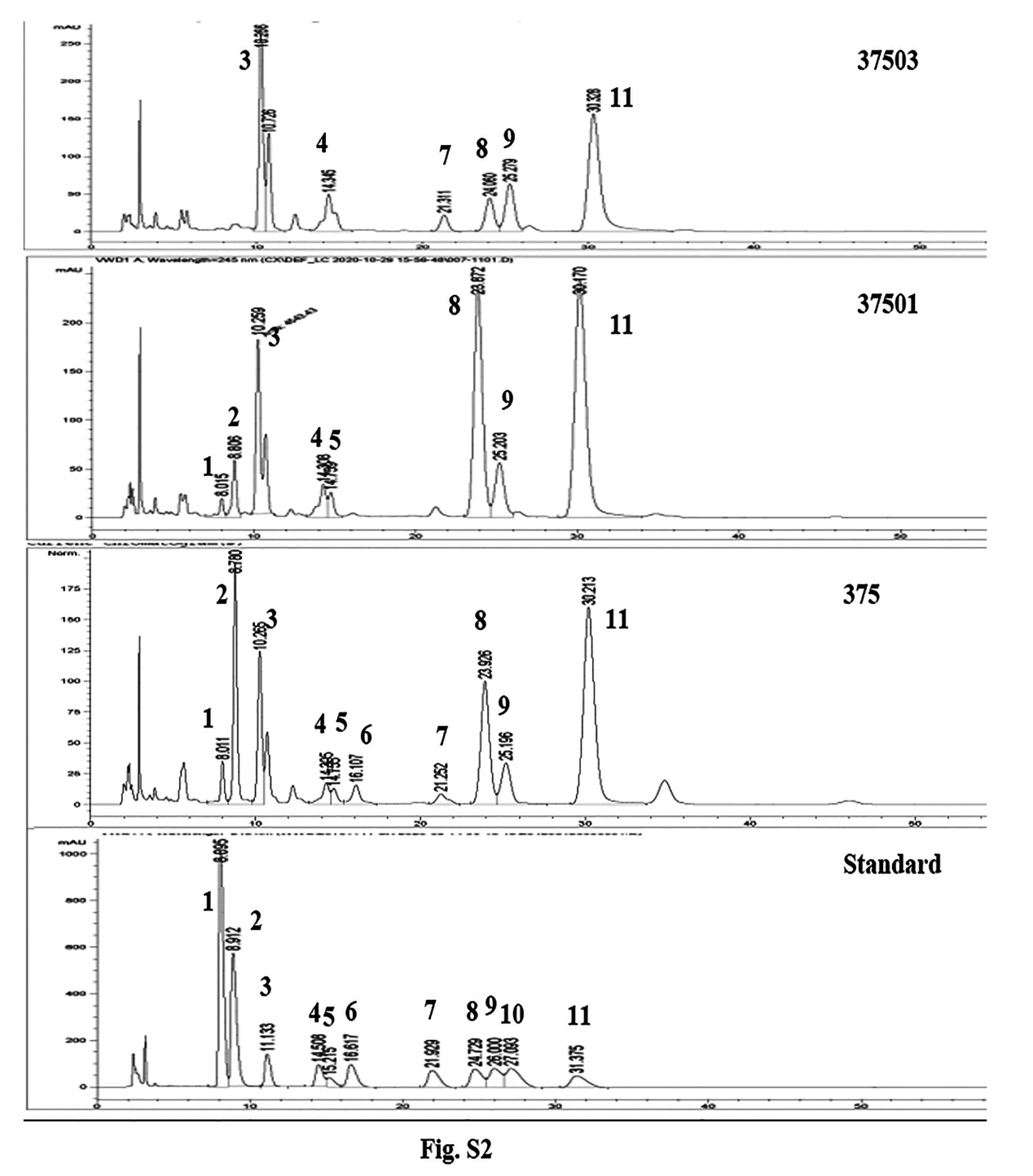

Fig. S2 Monosaccharide composition of polysaccharides 375, 37501 and 37503 according to standards (1, guluronic acid; 2, mannuronic acid; 3, Mannose; 4, Rhamnose; 5, Glucouronic acid; 6, Glactouronic acid; 7, Glucose; 8, Galactose; 9, Xylose; 10, Arabinose; 11, Fucose)

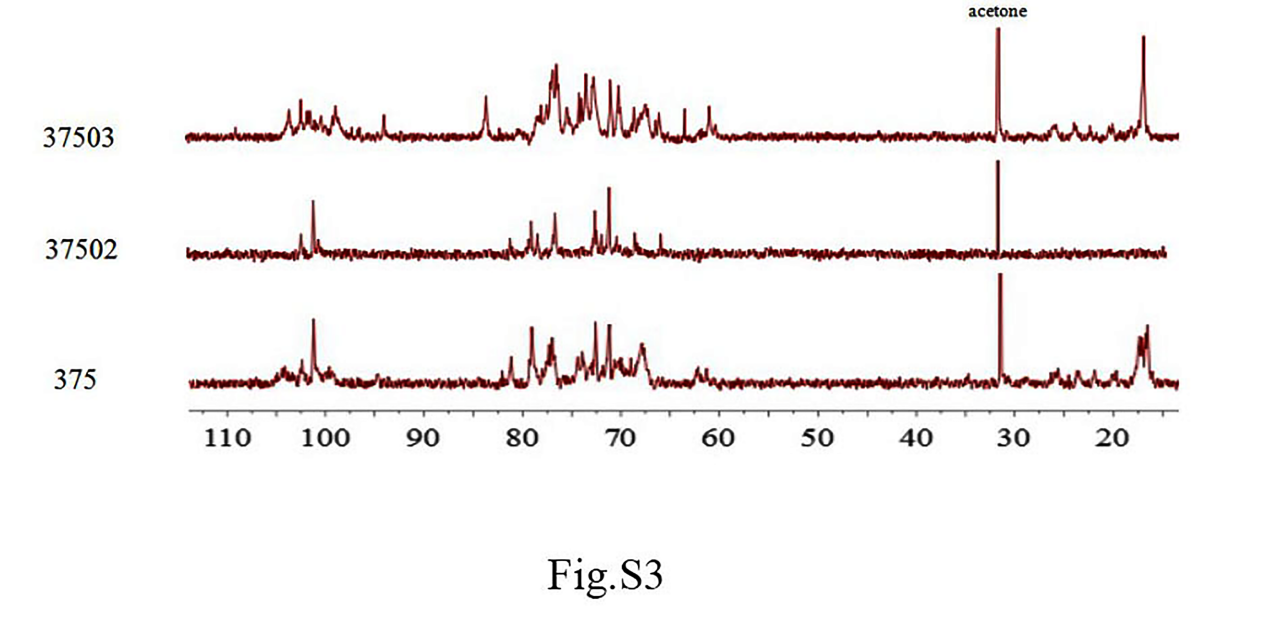

Fig. S3 ^13^C NMR spectra of polysaccharides 375, 37502 and 37503

**Acute toxic effects of mixture polysaccharides 375 on mice**

The objective of these studies was to perform a comparative evaluation of the acute toxicity of polysaccharides 375 in Balb/c mice.

[**Method**] Male C57BL/6J mice ages 8 weeks were purchased from Beijing Huafukang Laboratory Animal Co., Ltd. (Beijing, China). They were maintained under 12-h light/dark cycles and allowed ad libitum food and water. Animals and study protocol were approved by the Animal Care and Use Committee of Shanghai Institute of Materia Medica.

Mice (5 male, 5 female) were randomly divided into groups and intravenously administered with 0.9% saline solutions and different dosages of polysaccharides 375 (79.5 mg/kg, 106 mg/kg, 159 mg/kg and 238.5 mg/kg) for single dose, the median lethal dose (LD_50_) and 95% confidential intervals were measured. General behavior, adverse effects mortality and body weight were recorded for 14 days post treatment.

[**Result**] After the statistical analysis, the LD_50_ of polysaccharides 375 on mice was 136 mg/kg (95% confidence interval 115.128 mg/kg-164.187 mg/kg). Polysaccharides 375 significant toxicological changes occurred at the dose of 106 mg/kg, but not at the doses of 79.5 mg/kg. The results showed that at doses of 159 mg/kg and 238.5 mg/kg died within the first 2 days post-treatment. A 30% mortality rate was induced by 106 mg/kg, 60% mortality rate was induced by 159 mg/kg, and 100% mortality by 238.5mg/kg. The body weight gain of mice was not significantly changed at 79.5 mg/kg dosage of polysaccharides 375, but significantly changed at 106 mg/kg dosage of polysaccharides 375.

[**Conclusion**] Mixture polysaccharides 375 had certain acute toxic to mice. The results from mice showed that polysaccharides 375 could not induce any mortality at doses of 79.5 mg/kg, However, a 30% mortality rate was induced at 106 mg/kg of 375 and 60% mortality rate was induced at 159 mg/kg, while 100% mortality was at 238.5mg/kg.

Table S1. Acute toxicity evaluation on polysaccharide 375 by single dose intravenous administration on Balb/c mice

| does |  | Animal number |  | Death number |
| --- | --- | --- | --- | --- |
| 79. g/kg |  | 10 |  | 0 |
| 106.5 mg/kg |  | 10 |  | 3 |
| 159 mg/kg |  | 10 |  | 6 |
| 238.5 mg/kg |  | 10 |  | 10 |
